# Uncovering hidden biological processes by probabilistic filtering of single-cell data

**DOI:** 10.1101/2023.01.18.524512

**Authors:** Zoe Piran, Mor Nitzan

## Abstract

Elucidating underlying biological processes in single-cell data is an ongoing challenge and the number of methods that recapitulate dominant signals in such data has increased significantly. However, cellular populations encode multiple biological attributes, related to their spatial configuration, temporal trajectories, cell-cell interactions, and responses to environmental cues, which may be overshadowed by the dominant signal and thus much harder to recover. To approach this task, we developed SiFT (SIgnal FilTering), a method for filtering biological signals in single-cell data, thus uncovering underlying processes of interest. Utilizing existing prior knowledge and reconstruction tools for a specific biological signal, such as spatial structure, SiFT filters the signal and uncovers additional biological attributes. SiFT is applicable to a wide range of tasks, from the removal of unwanted variation in the data as a pre-processing step to revealing hidden biological structures. Applied for pre-processing, SiFT outperforms state-of-the-art methods for the removal of nuisance signals and cell cycle effects. To recover underlying biological structure, we use existing prior knowledge regarding liver zonation to filter the spatial *signal* from single-cell liver data thereby enhancing the temporal circadian signal the cells are encoding. Lastly, we showcase the applicability of SiFT in the case-control setting for studying COVID-19 disease. Filtering the healthy *signal*, based on reference samples from healthy donors, exposes disease-related dynamics in COVID-19 data and highlights disease informative cells and their underlying disease response pathways.

## Introduction

Cells encode information in their gene expression profiles about different facets of their identity, such as their spatial location within tissues, cell cycle phase, and disease stage. Recent years have seen a surge in computational methods for the reconstruction of such cellular facets from single-cell RNA-sequencing (scRNA-seq) data^1^. While these reconstruction methods were proven successful for the recovery of diverse signals, including spatial^2–4^ and temporal^5,6^ signals, the vast majority of these methods focus on reconstructing a single *signal* in the data, relying either on its dominance or based on sufficient prior knowledge (such as known marker genes). However, since cells encode multiple signals about their intrinsic state and extrinsic environment, focusing on a single *signal* (measured or recovered) is insufficient and may miss key cellular attributes. For example, while the cells’ spatial organization may be the dominant *signal* in a single-cell dataset, it may overshadow the temporal regulation of interest. In such a scenario, information about the reconstructed (e.g. spatial) signal can be utilized to filter it from the data and reveal further complex hidden attributes. Another example involves case-control comparisons, where information about healthy controls can be filtered from cells sampled from patients along disease progression or along response to treatment to uncover the subpopulation of cells affected by the disease and/or treatment and characterize their response.

The *signal* filtering approach was mainly explored for removing unwanted sources of variation as a pre-processing step. For example, multiple computational methods have been proposed for data integration and the removal of batch effects (e.g. bbknn^7^, Harmony^8^, and scVI^9^), or specifically for removing the cell cycle *signal*, a major source of bias in single-cell data^10–13^. This bias introduces large within-cell-type gene expression heterogeneity that can obscure the differences between cell types, which can in turn resurface once the cell cycle signal is filtered out. Yet, these methods are task-specific; cell cycle filtering approaches tend to account for known informative genes^12,13^ or take advantage of the *signal’s* cyclic structure^10^, while data integration methods (apart from scVI) typically focus on (a single) categorical factor to encode the different groups (batches) in the data. In addition, most integration methods, including scVI, provide a correction at the level of a joint latent representation of the cells, and not for the original count matrix or for individual genes, thus limiting the applicability of existing downstream analysis tools and requiring the development of case-specific methods^14^. Hence, altogether, the above methods cannot be used to filter out a generic biological *signal* from single-cell data.

To this end, Satija et al.^15^ suggested removing technical and cell cycle effects using a linear regression model. In this setting, independent linear models are fitted to predict gene expression with respect to a set of predefined variables. Then, for each variable independently, the fitted linear model is regressed from each gene. However, this strategy is restricted to the fit of the linear model and does not allow for additional user inputs to adjust the removal process to ensure that desired biological components are not removed from the data.

Here we introduce SiFT (***Si**gnal **F**il**t**ering*), a diverse and robust framework for filtering signals induced by different biological processes in single-cell data, thus uncovering underlying processes of interest. To do so, we compute a probabilistic cell-cell similarity kernel, which captures the similarity between cells according to the biological *signal* we wish to filter. Using this kernel, we obtain a projection of the cells onto the *signal* in gene expression space. By deducting this projection from the original data, we filter the *signal*-related information and uncover additional, *hidden* cellular attributes.

We begin by demonstrating that SiFT can successfully and efficiently remove sources of unwanted variation in the data while preserving biological attributes. This is showcased by applying SiFT to filter nuisance signals in Drosophila wing disc development single-cell data, and removing cell cycle effects from a semi-synthetic single-cell dataset mimicking the existence of two sub-clones. SiFT outperforms state-of-the-art methods in all cases. Next, we exemplify SiFT’s ability to expose and enhance underlying biological signals. To do so we use prior knowledge regarding liver zonation to filter the spatial signal from single-cell liver data, thereby enhancing the temporal circadian signal encoded by the cells. Finally, we demonstrate the applicability of SiFT in the case-control setting. We apply SiFT to COVID-19 dataset, using healthy samples as a reference we filter the joint healthy *signal* and expose disease-related dynamics.

SiFT is available as a scalable, user-friendly open-source software package https://github.com/nitzanlab/sift-sc, along with documentation and tutorials at https://sift-sc.readthedocs.io.

## Results

### Revealing hidden biological signals using SiFT

The SiFT framework leverages known relationships between cells to expose additional, underlying structures in single-cell data (Figure 1). Consider the scenario where each cell has two attributes, which we will term here *shape* and *color*. Now, assume that we have experimentally measured, or we can computationally recover the *color* identity (e.g. by coupling known marker genes to a clustering of the data, Figure 1(b)). Yet, the biological signal which remains meaningful to uncover is the *shape* of the cell, for which no prior knowledge exists. By using SiFT to remove the *color* signal from the data, we will be able to uncover the *shape* signal.

**Figure 1:**
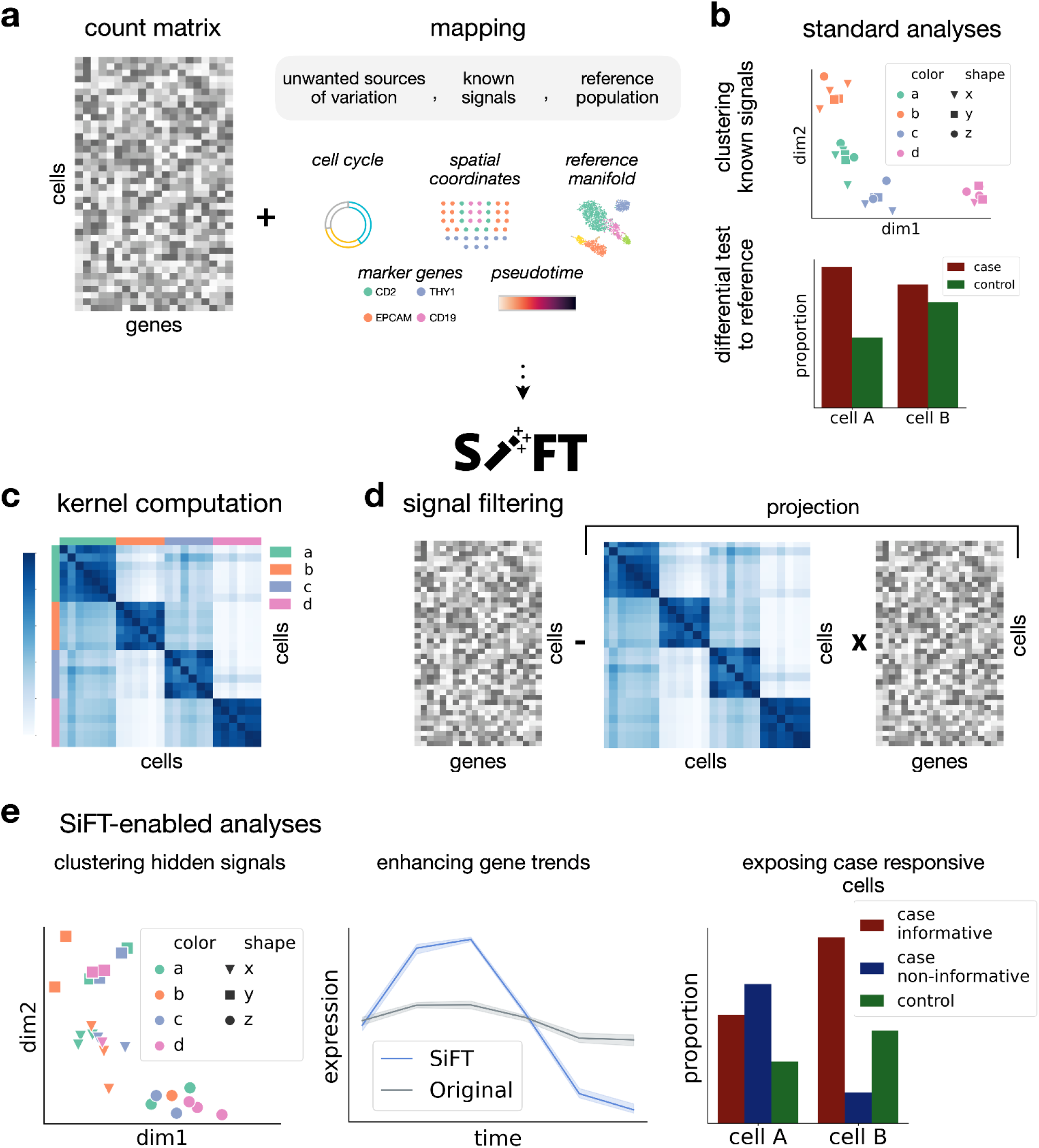
Overview of the SiFT algorithm. **(a)** SiFT takes as input a count matrix and a pre-computed mapping of the cells. The mapping can be either continuous or discrete and either univariate or multidimensional. **(b)** Standard analysis recovers dominant *signals* in the data. A comparative abundance test is performed between case and control; without correcting for case-responsive cells, *cells of type A* are found to be more prevalent in disease state and hence will typically be classified as disease informative **(c)-(d)** The *SiFT* pipeline. **(c)** A cell-cell similarity kernel is computed based on the given signal mapping. **(d)** Filtering is performed by projecting the kernel onto the count matrix and deducing the projection from the original count matrix. **(e)** After filtering, the hidden structure is exposed and easily recovered in downstream analysis. SiFT allows labeling according to hidden signals (left), enhances underlying gene trends (middle), and uses a reference control population to identify cells that are responsive for the case, correcting the naive abundance test and identifying that *cells of type B* contain a larger population of cells informative of disease state (right).

SiFT takes as input both an expression count matrix, as well as knowledge about a specific *signal* encoded by attributes of the cells. The latter can be provided as a mapping of the cells using deterministic labels (e.g. cell cycle stage or spatial coordinates), a set of marker genes, pseudotime ordering, or a latent space representation of the cells (see Methods, Figure 1(a)). These attributes are used to compute a cell-cell similarity kernel with respect to the encoded *signal*. Alternatively, the mapping to the *signal* can be based on a population of reference cells (e.g. control cells in a case-control setting). Then, the cell-cell similarity kernel is computed only with respect to the reference cells to capture the reference (e.g. control) *signal*.

In general, the kernel captures distances between cells in the *signal* space, thus encoding the cells’ similarity with respect to the signal to be filtered. We define three main variants of cell-cell similarity kernels: a *mapping, k*-nearest-neighbo*r* (*knn*), and a *distance* kernel. The choice of the kernel type relies on the structure and knowledge regarding the existing signal attribute, where the kernels mainly differ in the type of prior knowledge they require over the attributes (for further discussion see Methods). The *mapping kernel* relies on a stochastic or deterministic mapping of the cells to a given domain. Such mapping results in cell labels, including cell-type labels or temporal labels generated by binning of a pseudotime trajectory. The *knn* and *distance kernels* rely on a joint space representation of the cells over which a corresponding distance metric can be defined. Such spaces can be generated, for example, by restricting the original single-cell data to a set of marker genes, or a latent space representation of the cells based on single-cell variational inference (scVI)^9,16^. Alternatively, the user can supply a pre-computed cell-cell similarity kernel that captures information they aim to filter.

Given a cell-cell similarity kernel, we obtain a projection of the expression count matrix onto the signal we seek to filter using matrix multiplication (see Methods). With this construction, the projection can be interpreted as the portion of the expression associated with the known signal. Hence, we can deduce it from the original count matrix, and obtain a filtered representation of the data (Figure 1(d)). We can now utilize various analysis tools to study the filtered data and recover the underlying biological signals it encodes (Figure 1(e)).

The filtering of the kernel’s projection over the gene expression removes effects induced by the original signal. Consequently, the remaining, filtered gene expression stands for the deviation of each cell’s gene expression from the expected, or typical expression in its neighboring cells. This deviation captures the attributes associated with the additional biological signals in the data. Thereby SiFT provides a corrected count matrix (which can contain negative values or be corrected by a pseudo count, see Methods) encoding the contribution from underlying processes, making it easier to study them.

### SIFT efficiently removes unwanted sources of variation

Experimental data often contains unwanted sources of variation which obscure the biological signal of interest. These can be discrete (e.g. sex label) or continuous (e.g cell cycle phase) signals. A desired pre-processing step is to remove these contributions. An optimal removal procedure is one that efficiently removes the unwanted signal while preserving the biological attributes, is generic in the sense that it can be adapted to diverse settings, and can easily be included in the analysis pipeline, in terms of implementation and scalability with respect to dataset size. We turn to show that SiFT meets all of these criteria.

We start with a single-cell transcriptomics dataset of the Drosophila wing disc which was shown to suffer from unwanted sources of variation due to cell cycle and sex signals^17^. The original embedding of the cells reflects the bias induced by both the cell cycle and sex signals within each batch (Figure 2(a)). Using SiFT we can filter these signals and uncover the underlying biologically meaningful variability. As input to SiFT, we use the measured gene expression and a set of marker genes that captures the cell cycle and sex-related variation (see Supplementary table 1). We define a *knn* kernel using the marker genes, meaning that we identified the neighborhood of the cells using a similarity measure based on cell cycle and sex genes (Supplementary Figure 1). Then, this kernel is used to filter out the unwanted signals.

**Figure 2:**
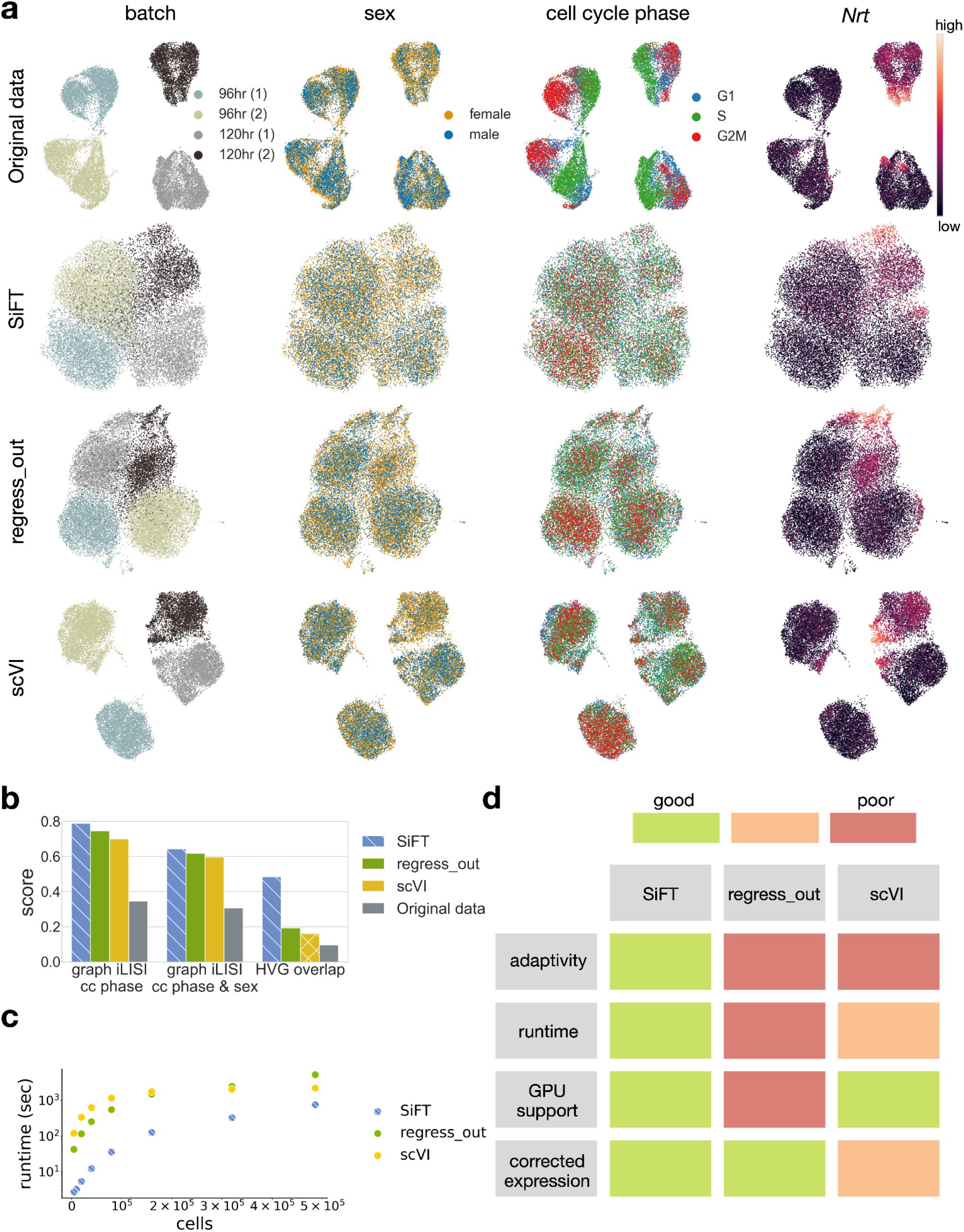
SiFT is optimal for removing unwanted variation from single-cell data. **(a)-(b)** transcriptomics of the Drosophila wing disc development (data: Everetts et al.^17^). **(a)** UMAP embeddings^20^ following different data correction procedures (rows) and colored by different covariates of unwanted sources of variation (columns). Rows (top to bottom) show uncorrected data (Original data), SiFT filtering using *knn* kernel (SiFT), Scanpy’s “scanpy.pp.regress_out()” (regress_out), and scVI latent space with continuous covariates (scVI). Columns (left to right) show the cells colored by batch label, sex label, cell cycle phase, and Nrt, a novel Hedgehog target gene identified by^17^. **(b)** Integration and biological preservation scores per method. Scores (left to right): graph iLISI score evaluated for cell cycle phase label, graph iLISI score evaluated for cell cycle phase and sex label, and overlap of the highly variable genes with a set of genes of biological interest (see Methods). **(c)** The runtime of the methods on subsampled versions of the Heart Cell Atlas dataset^21^. **(d)** A table summarizing several criteria regarding the different methods: (top to bottom) *adaptive*; relates to the flexibility of the method, support of different filtering procedures allowing for optimization of the task. Both of the compared methods, regress_out and scVI, do not support any additional parameters apart from the variable of interest. *runtime*; color indicates the overall scalability of the method. A combined measurement of overall runtime and scalability across magnitudes, as depicted in (c). *GPU support*; implemented support of GPU acceleration. *corrected expression*; indicates whether the method outputs the corrected gene expression (for a disclaimer regarding scVI’s applicability to this evaluation as it requires imputation of the corrected gene expression see Methods).

The SiFT-corrected embedding of the cells shows a homogeneous representation with respect to the sex and cell cycle phase signals, which we aimed to filter (Figure 2(a)). That is, in contrast to the original data, labels are not visibly separable in the latent space representation (Figure 2(a)), and the marker genes’ spatial gradients are removed (Supplementary Figure 2). This qualitative result is supported quantitatively by the graph iLISI score with respect to both cell cycle labels as well as cell cycle and sex labels (see Methods), which shows that SiFT can successfully remove unwanted variation in single-cell data (Figure 2(b)).

Furthermore, SiFT outperforms available baselines in this task, including Scanpy’s regress out function^18^ (a python implementation of the linear regression suggested by Satija et al.^15^) (regress_out) and scVI conditioned on the expression of the set of nuisance genes^9^ (scVI). This can be seen by qualitatively comparing the labels’ separation in the latent space representation (Figure 2(a)), and quantitatively with respect to the graph iLISI score (Figure 2(b)).

Importantly, while filtering unwanted variance in the data SIFT preserves biological variation that was not targeted for filtering, as 50% of the genes of interest (including genes reported in the original analysis of this data^17^ and in^19^, see Supplementary table 1) present in the corrected highly variable gene set (see Methods, Figure 2(b)), while regress_out captures only 20% (see Methods). Such preservation of biological variation is visualized for Neurotactin (*Nrt*) and midline (mid), downstream Hedgehog pathway targets in the adult muscle precursors, and patched (*ptc*), a receptor of the Hedgehog ligand (Figure 2(a), Supplementary Figure 2). When coupled to standard batch integration methods (e.g. Harmony^8^ or bbknn^7^), SiFT attains better performance in the complete integration task when compared to regress_out followed by Harmony or bbknn, or scVI applied with batch correction with continuous covariates (Supplementary Figure 3).

Next, we turn to show that SiFT is scalable and can be applied to large single-cell datasets. Hence, we benchmark runtime performance on the Heart Cell Atlas dataset, composed of nearly 500,000 cells^21^. To define a filtering task we add random features in the form of continuous random noise. The features are used as the mapping for the signal we wish to remove (see Methods). Given this task we test the runtime of the removal algorithms (SiFT, scVI, and regress_out) over increasing subsamples of the data (Figure 2(c)). This showcases SiFT’s efficiency and scalability to large single-cell datasets, with a runtime of 12 min, 36 min, and 1 h 26 for SiFT, scVI, and regress_out, respectively over the complete dataset.

Together, SiFT meets all criteria desired for the successful filtering of unwanted *signals* in single-cell data (Figure 2(d)); Its versatility in kernel computation allows adapting the filtering to uncover and preserve the biological signal of interest, its runtime efficiency along with GPU support allows scaling to large datasets, and since filtering is performed directly on the input count matrix, it can naturally be incorporated into the data pre-processing step and followed by any downstream analysis procedure.

### SiFT exposes underlying biological heterogeneity

An important aspect of the filtering procedure is the ability to expose underlying biological attributes. The cell cycle introduces heterogeneity that can obscure other biological differences between cells, and so removing its effect improves the inference of inherent biological diversity^11,13^. Hence, many dedicated methods were introduced for this task, amongst them are Cyclum^10^, an autoencoder-based approach for identifying circular trajectories in gene expression space, Seurat^12^, which uses a linear model to find the relationship between gene expression levels and marker genes scores it assigns to each cell, and ccRemover^13^, a PCA-based method that identifies and then removes components related to the cell cycle.

We consider a synthetic manipulation of single-cell data curated by^10^ which consists of two sub-clones and show that SiFT can successfully remove the cell cycle effect and enhance the sub-clone separability in the data. The two sub-clones provide a supervised setting resembling a biological signal we intend to preserve and expose. We used mouse embryonic stem cell (mESC) data as one clone^22^ and a second clone was created by doubling the expression levels of a randomly selected set of genes containing variable numbers of cell cycle and non cell cycle related genes. The cell cycle stage can be easily recovered in the original data, but not in the sub-clone where it is hidden (Figure 3(a)).

**Figure 3:**
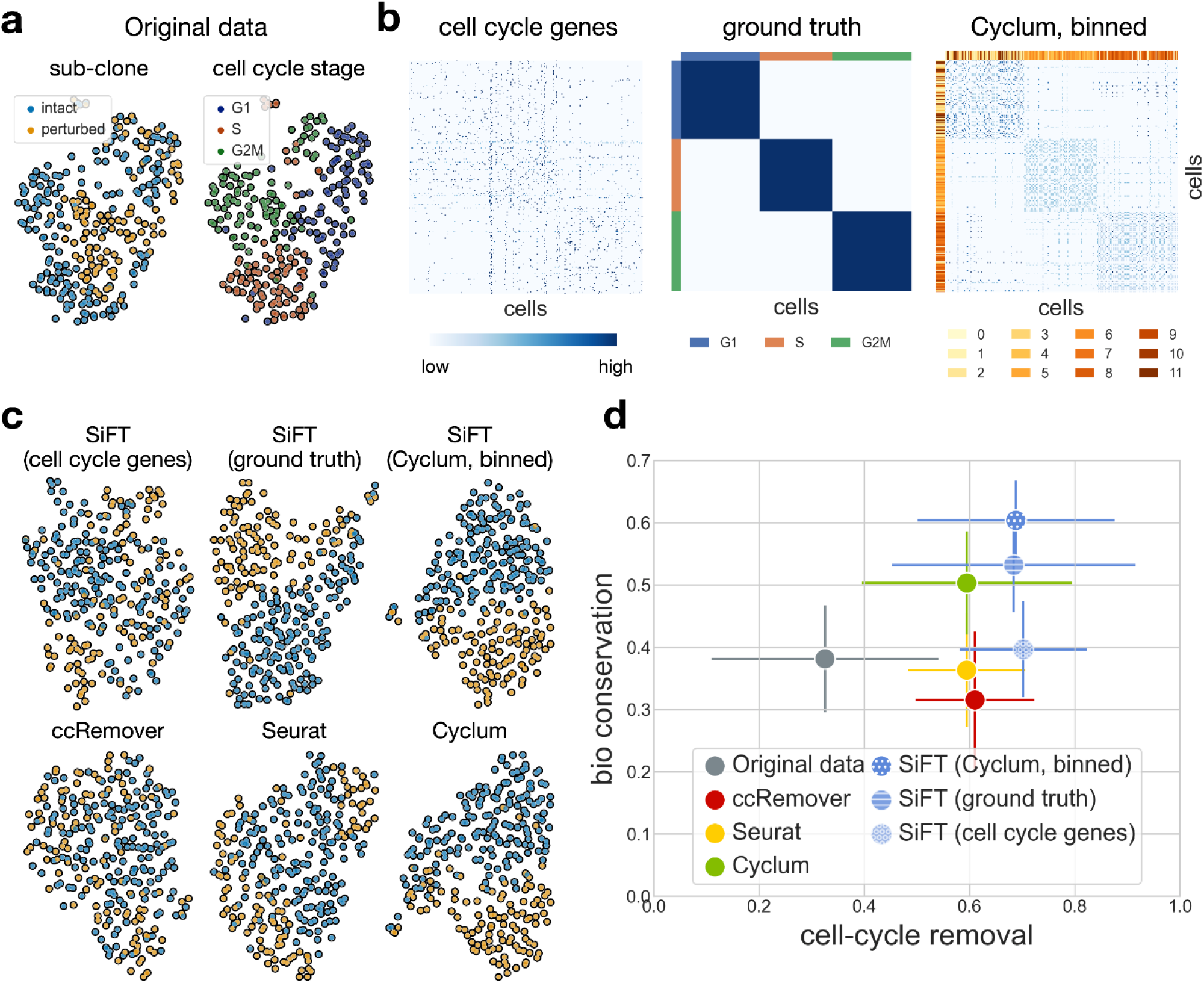
Filtering the cell cycle effects from the virtual tumor data consisting of two sub-clones. **(a)** A UMAP of the original data colored by sub-clone identity (left) and cell cycle stage labels (right). **(b)** The different cell-cell similarity kernels as defined by SiFT, cells are ordered according to the ground truth cell cycle stage. (left) a *knn* kernel, neighbors defined using the gene expression of a set of cell cycle marker genes. (center) a *mapping kernel* based on the ground truth cell cycle stage, the row(col) colors depict the cells’ label. (right) a *mapping kernel* based on binning of the Cyclum pseudotime, the row(col) colors depicts the cells’ bin. **(c)** UMAP of the filtered data colored by the sub-clone identity. The top row presents SiFT filtered data, (left) cell cycle marker genes (center) ground truth labels (right) Cyclum pseudotime binning Bottom row presents compared methods, (left) ccRemover (center) Seurat (right) Cyclum. **(d)** Scatter plot of the mean overall *bio conservation score* against mean overall *cell cycle removal scores* using the metrics defined in ^23^ (see Methods). The error bars indicate the mean standard error.

To quantify the performance and assess the separability of the sub-clones along with the mixing of the cell cycle stages, we utilize a set of metrics for integration accuracy^23^. The metrics are divided into two categories, removal of batch effects and conservation of biological variance (all scores are scaled between 0 to 1), where the cell cycle stage labels are considered as the batches in the data. Hence in our context, the batch effects removal scores correspond to the cell cycle stages mixing. Similarly, the cell subclone identity label is used as the anchor for the preservation of biological signals, thus used in the metrics for biological variance (see Methods).

We start with an independent evaluation of SiFT using only a set of known cell cycle marker genes as input to compute a *knn* kernel (see Methods, Figure 3(b)). The marker genes define the manifold for the computation of the neighbors. Here, the low dimensional representation of the cells does not expose clear visual separation between the sub-clones (Figure 3(c)), yet the quantitative assessment ensures it performs well for the desired task and more specifically outperforms Seurat which relies on the same prior knowledge (Figure 3(d)). Next, we show how additional prior knowledge can enhance the accuracy of SiFT by two alternative mappings for the cell cycle signal. The ground truth cell cycle stage (provided with the data), and the binned representation of the pseudotime inferred by Cyclum^10^. Both are then used to construct mapping kernels. The mapping kernels showcase a block structure depicting the three cell cycle stages while the *knn* kernel captures more subtle relations between the cells (Figure 3(b)).

Following filtering of cell cycle effects, the visual separability of the sub-clones is substantially enhanced by SiFT, as well as by Cyclum, using either the ground truth labels or Cyclum’s pseudotime (Figure 3(c)). SiFT consistently attains quantitatively higher scores in cell cycle removal metrics in comparison to baseline methods, and higher scores in biological conservation when relying either on ground truth or Cyclum based mapping to be filtered (Figure 3(d)).

### Filtering spatial signals recovers temporal information

Cellular gene expression is regulated by spatial and temporal signals, posing a challenge of decoupling and studying the interplay between the two. The liver stands as an example of a tissue undergoing strong spatial and temporal regulation; it consists of repeating anatomical units termed liver lobules, and sub-lobule zones performing distinct functions. Liver zonation refers to functions that are non-uniformly distributed along the lobule radial axis. Beyond the spatial structure, the liver is also subject to temporal regulation, consisting of the circadian clock, systemic signals, and feeding rhythms^24,25^. While the liver zonation *signal* has been intensively studied^25,26^, less is known regarding the temporal signal^24^. We turn to demonstrate how knowledge regarding the spatial zonation *signal* can be used by SiFT to enhance the circadian clock trajectory. We consider a dataset from^24^, labeled for temporal processes, as data were collected at four different equally-spaced time points along the day, and with known marker genes for the spatial zonation signal^26^ (see Supplementary Table 2). Hence, this data allows us to computationally recover the spatial structure for filtering (Figure 4(a)).

**Figure 4:**
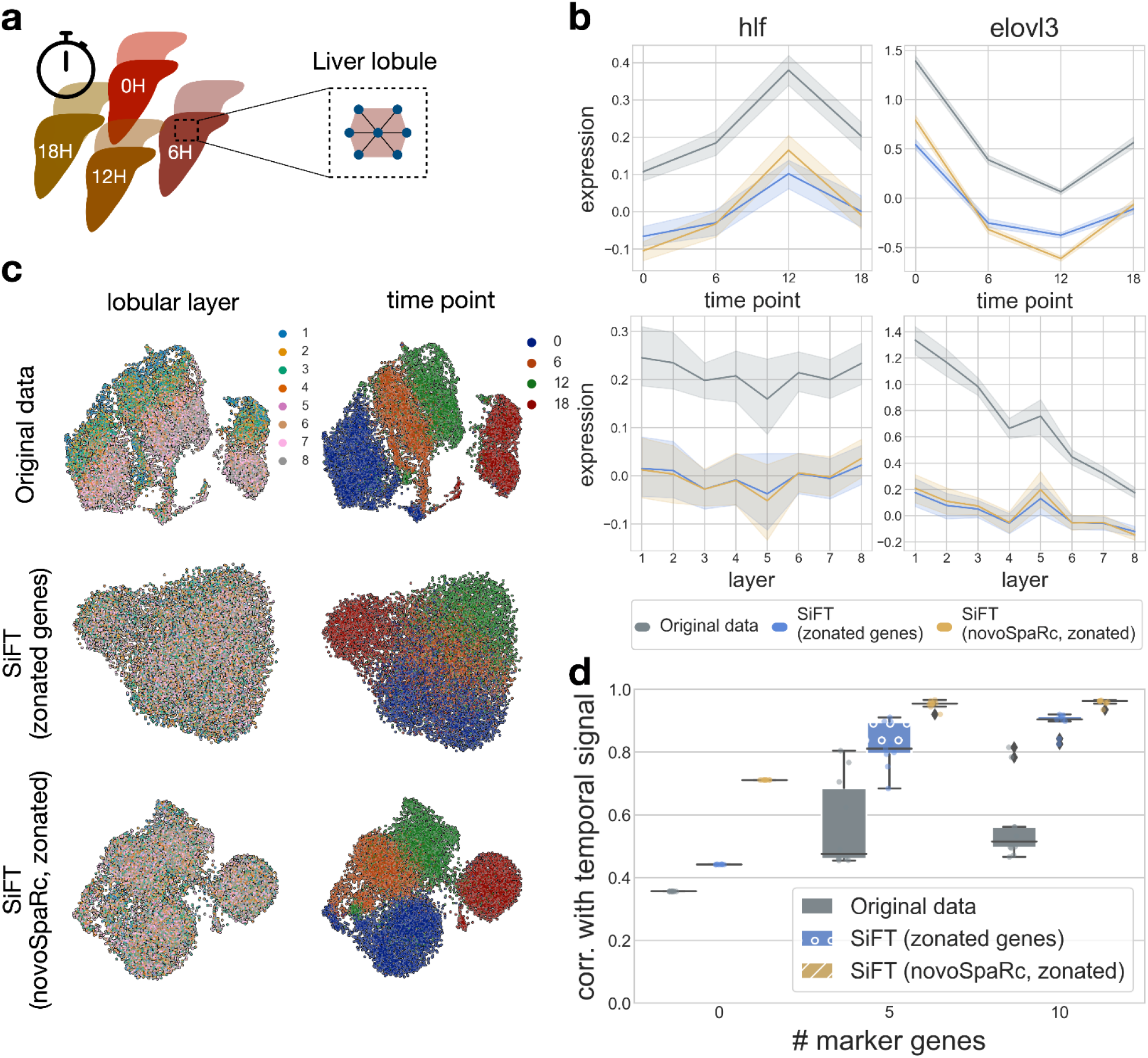
Enhancing circadian clock signal in the mammalian liver by filtering the spatial zonation signals using a mapping to lobular layers. **(a)** Single-cell RNA-seq of hepatocytes isolated at four different times within a day ^24^ **(b)** Reconstructed temporal (top) and spatial (bottom) expression patterns of zonated and rhythmic genes (*hlf* and *elovl3*) **(c)** A UMAP representation of the original data (top), SiFT filtered data based on zonated genes (middle), and novoSpaRc zonated mapping (bottom). Plots are colored by the lobular layer (left) and time point (right). **(d)** Correlation between reconstructed and original rhythmic signals as a function of the number of temporal reference genes used for the reconstruction.

While SiFT can take as input the set of spatial marker genes directly, it can be beneficial to use existing methods dedicated to the spatial reconstruction and provide the spatial mapping directly as input to SiFT. Using SiFT solely with marker genes exposes the temporal signal and by providing the spatial reconstruction input, we can further improve its performance. To obtain a spatial reconstruction we use novoSpaRc^2,3^, providing a mapping of the spatial organization of the cells. NovoSpaRc is an optimal transport based method for manifold mapping of scRNA-seq data which can take as input prior knowledge in the form of marker genes (see Methods). The mapping, as inferred by novoSpaRc, to an eight-layer tissue representing the lobular liver layers, is used by SiFT to construct a *mapping kernel* (Supplementary Figure 5). The set of spatial marker genes is used by SiFT to define a *knn kernel*, where distances between the cells are computed based on the similarities in the spatially zonated genes (Supplementary Figure 5).

In both settings, using zonated genes or relying on novoSpaRc mapping, applying SiFT successfully removes the zonation signature yet preserved the visual separability based on the circadian trajectory over a UMAP representation of the data (Figure 4(c)). This is further supported by comparing the expression of two zonated and rhythmic genes (*hlf* and *elovl3*) before and after the application of SiFT (Figure 4(b)): The genes’ temporal trend is preserved (Figure 4(b), top row), yet the spatial variation is eliminated (Figure 4(b), bottom row).

We look to quantitatively test the performance by assessing the correlation of the reconstruction with the reported temporal trajectory as a function of the number of temporally informative genes (marker genes used for the reconstruction, see Supplementary Table 2). To do so, we utilize novoSpaRc again, this time to recover the temporal trajectory by mapping to four cyclic locations. We find that the quality of temporal trajectory reconstruction with SiFT (using novoSpaRc, zonated), even without any temporal reference, is better than the reconstruction based on the original data with 10 marker genes (SiFT (0 markers) = 0.71 ± 0.0, Original (10 markers) 0.56 ± 0.1), implying that by utilizing SiFT, prior knowledge of the zonation signature exposes the underlying circadian clock trajectory and provides sufficient information for its successful recovery (correlation of 0.71). Further, even in the simpler setting, applying SiFT using zonated genes, improves the quality of temporal reconstruction without any marker genes (0.44 ± 0.0). At last, In both of the SiFT settings, the reconstruction quality substantially improves upon the addition of informative genes and reaches near-perfect performance using 10 markers (SiFT (zonated genes) = 0.89 ± 0.3, SiFT (novoSpaRc, zonated) = 0.96 ± 0.0, Original 0.56 ± 0.1) (Figure 4(d)).

### Using a reference of healthy control cells to recover disease signature

An essential step in studying disease is decoupling disease signatures from the healthy state by characterization of the healthy *signal* in clinical samples. SiFT can be used for this task, recovering disease-specific signatures in individual cells, by *filtering* the healthy trajectory based on healthy control samples. This allows for enhancing the disease signature, identifying cells that are more informative for disease states, and studying the different types of disease response pathways (Figure 1). Of note, existing analysis pipelines approach this task by integrating all samples, from both healthy controls and disease patients, and then performing comparative analysis, for example, by performing differential expression analysis between common cell types or assessing differences in cellular composition between health and disease^27–30^. The extent of this type of analysis is limited as it relies on bulk comparison and does not expose the disease trajectory in individual cells.

We consider single-cell transcriptomes from peripheral blood mononuclear cells (PBMCs) from individuals with asymptomatic, mild, moderate, severe, and critical COVID-19 (*n* = 90 individuals) and controls (*n* = 23 individuals, Figure 5(a))^27^. We use a harmonized PCA space (as reported in Stephenson et al.^27^), and restrict the set of reference cells to the healthy control samples. We then define a knn kernel based on this space to compute the similarity between each cell from individuals with COVID-19 relative to the healthy population (see Methods). Thus, applying SiFT is expected to filter the signal of the healthy trajectory (as present in each cell independently) from the data and expose the disease response in each cell (Figure 5(b)).

**Figure 5:**
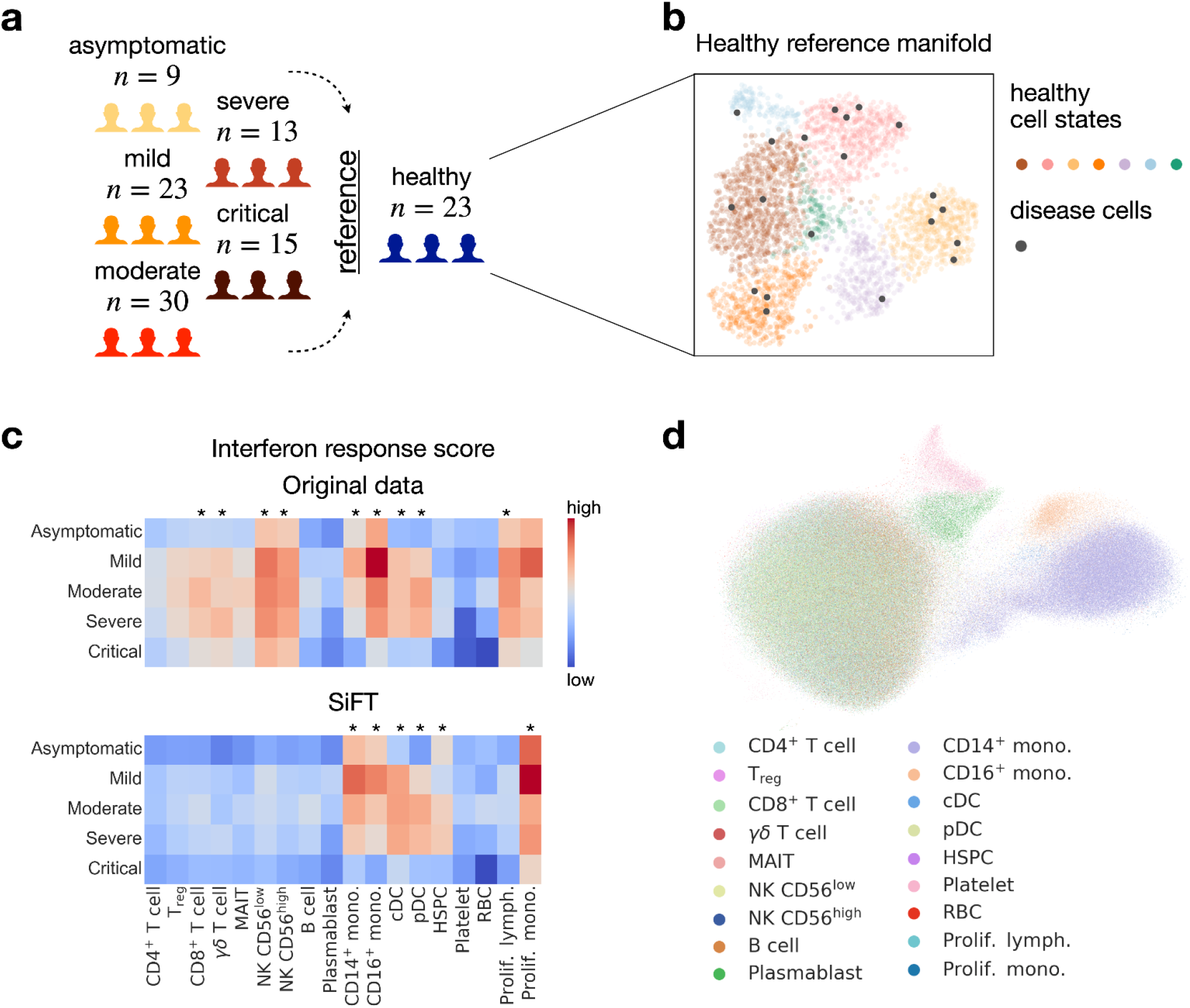
Exposing disease signature through the removal of similarity to healthy control cells in single-cell PBMCs transcriptomes from individuals with asymptomatic, mild, moderate, severe, and critical COVID-19. **(a)** Overview of the participants included and the data collected by Stephenson et al.^27^. A total of *n* = 90 COVID-19 patients and *n* = 23 control individuals. **(b)** Abstract illustration of the healthy reference mapping procedure. Cells from patient samples are projected onto the healthy reference manifold. **(c)** Enrichment of interferon response of each cell state separated by disease severity. Shown for original data (top) and SiFTed data (bottom). IFN response score was calculated using a published gene list (GO:0034340). Statistical tests were performed with a Mann-Whitney U test between the cell types. Cell types are considered statistically significant if *p_value_* < 0.05 (denoted by *). **(d)** UMAP visualizations of all cells in the SiFTed data colored by reported cell type.

Patients with COVID-19 present with an abnormal immune landscape, characterized by overactivated inflammatory, innate immune response, and impaired protective, adaptive immune response^31^. Recent studies revealed the dynamic changes in peripheral immune cells, both in transcriptional states and population size over the course of COVID-19^28,32^. The SiFTed representation recovered cell types involved in innate immune response, based on the interferon response (IFN) score (*monocytes, DCs*, and *HSPC*) whereas the same analysis over the original data failed to expose the relevant cell types ^27^ (Figure 5(c); statistical tests were performed with a Mann-Whitney U test between the cell types). While this acts as validation that SiFT can successfully recover results obtained from direct comparative analysis^27^, we next show how using SiFT can go beyond current analysis methods.

Considerable effort has been put into identifying the expansion of different cell types in response to COVID-19 infection^27,33^. This is assessed by comparing the cell type-specific population size between the disease and control samples. This analysis, however, is insufficient as it does not expose the extent to which these cells respond to the infection and furthermore, which fraction of the expanded population contains information regarding the disease state. SiFT allows for both as it refines and extends the initial cell type classification with respect to the disease response, hence exposing the distinct underlying disease signature and identifying the cells that are informative for the disease state. Intuitively, by SiFTing the healthy *signal* we expect that cells of a given cell type with a dominant disease response will preserve their identity (and will later be clustered together) and that remaining cells, with a less distinct response (more similar to healthy cells), will tend to cluster with non-informative cells, as most of their *signal* was removed. Indeed, the latent space representation of the SiFTed data exposes sub-populations of specific cell types (*CD*14^+^ *mono*., *CD*16^+^ *mono, Plasmablast*, and *Platelets*) while other cell types got mixed (Figure 5(d)). Importantly, the exposed cell types are known to undergo changes in response to COVID-19 infection^34^.

Next, to obtain a classification of cells as disease informative, we used a *cluster_purity* test over Leiden-based clusters in the SiFTed data with respect to cell type labels (see Methods, Supplementary Figure 6, Figure 6(a)). Under the *cluster_purity* test, a cluster is classified as disease informative if the fraction of the most prominent cell type in it exceeds a threshold value *τ* = 0.55. Gene-set enrichment analysis (GSEA) over the differentially expressed genes between the two groups (disease informative/disease non-informative) supported this classification and identified pathways associated with inflammation including response to virus, response to type I interferon, IFN*α* response, inflammatory response, and regulation of viral process in the disease informative cells (Figure 6(b)). Additionally, under the classification of disease informative cells, the IFN response signature was refined by enhancing the expected pattern of disease informative cells and exemplifying the lack of disease-related signal of remaining disease non-informative cells (Supplementary Figure 6). In accordance with this, the disease informative cells exhibited overexpression of type I/III interferon response-related genes (Supplementary Figure 6), which were recently reported in genome-wide association studies (GWAS) for COVID-19 susceptibility^35,36^.

**Figure 6:**
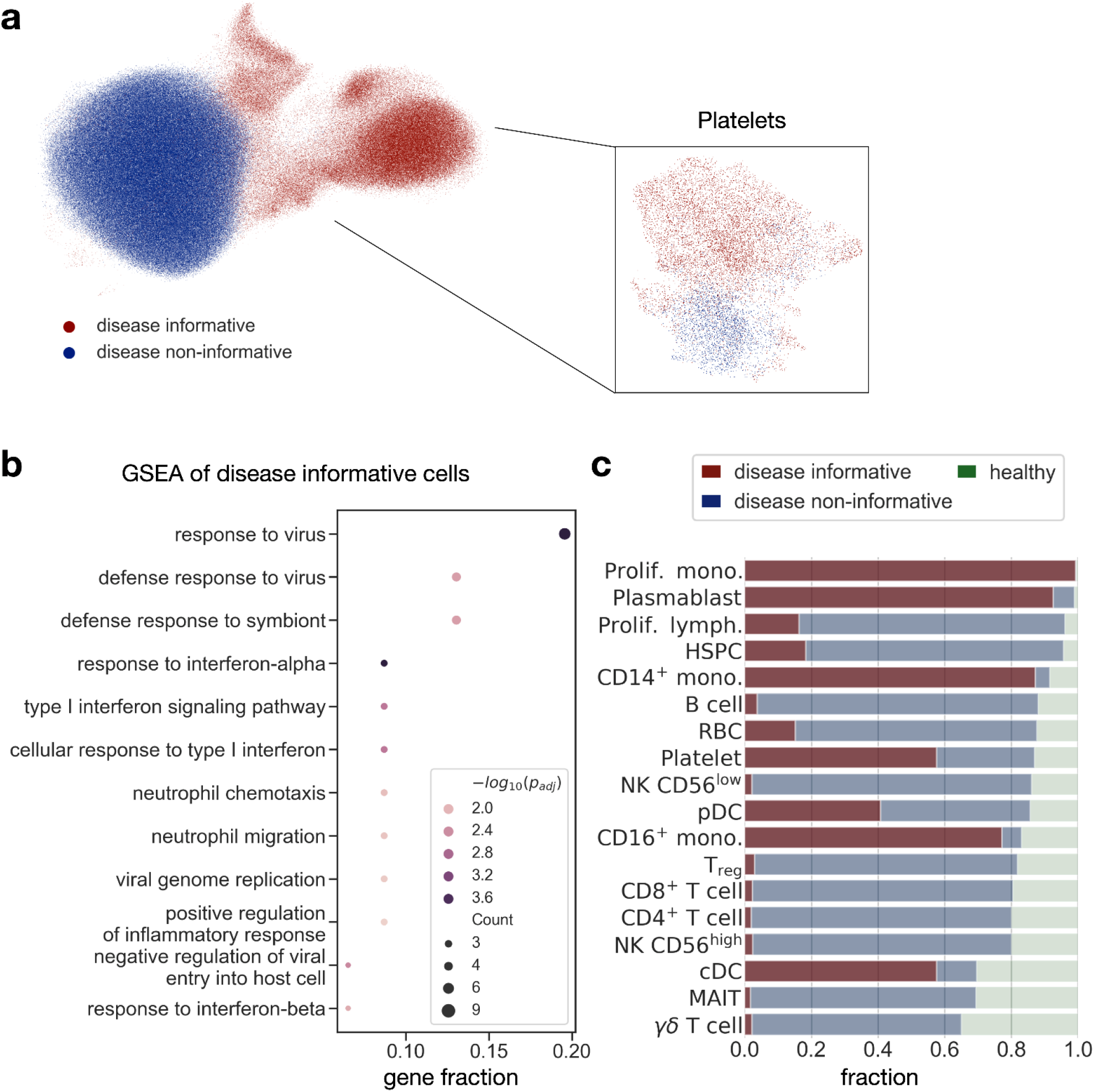
Classification of cells from clinical samples to disease informative (non-informative). **(a)** UMAP visualizations of all cells in the SiFTed data colored by association to disease informative (non-informative) clusters. Leiden clusters in the SiFTed data were classified according to *cluster_purity* score (see Methods). **(b)** GSEA of differentially expressed genes in the disease informative cluster (compared to non-informative, using top 50 genes). The size of the circles indicates the number of genes. Color indicates the magnitude of −log_10_(*p_adj_*). **(c)** Disease informative bar plot of the proportion of cell populations, separated into disease informative, disease non-informative, and healthy (according to the assignment shown in (a)). Cell types are sorted according to the fraction of disease cells.

The classification of disease informative/non-informative clusters provides a refinement to the cell type proportion analysis in the disease vs. healthy control (Figure 6(c)). Importantly, for a given cell type the fraction of inferred informative cells is not dictated by its prevalence in clinical samples. For example, *Prolif. lymph*. was ranked as the third most prevalent cell type in clinical samples (96% of the cells in disease state), yet, only 15% of *Prolif. lymph*. cells were associated with a disease informative cluster.

Furthermore, identifying disease informative and non-informative clusters exposes populations of interest and allows a focused analysis of the informative cells. For example, while increased platelet activation and coagulation abnormalities were previously reported in COVID-19 patients^27,37^, we identified two distinct subpopulations of *Platelets*: informative and non-informative for disease state (Figure 6(a)). A GSEA analysis based on differential gene expression between the disease informative and non-informative clusters identified pathways associated with coagulation, hemostasis, and antimicrobial humoral response in the disease informative cells, as well as increased expression of surface proteins indicative of platelet activation in the disease informative cells (Supplementary Figure 6).

## Discussion

We presented SiFT, a method aiming at discovering hidden cellular processes by filtering out a known or reconstructed *signal* from single-cell gene expression data. The SiFT procedure starts by defining a cell-cell similarity kernel, capturing similarities with respect to the *signal* to be filtered. This kernel is then used to obtain a projection of gene expression onto the *signal*, which is then removed from the original expression.

We have shown that filtering signals by SiFT can expose the underlying, biologically meaningful structure in the data over a wide range of tasks. First, we showcased its ability to successfully filter unwanted sources of variation caused by nuisance *signals* in the data while preserving biological signals of interest. When focusing on removing cell cycle effects, a major source of bias in single-cell data, in a semi-simulated setting we showed that SiFT outperforms state-of-the-art methods for the task. A substantial advantage of SiFT is its ability to use existing prior knowledge to reveal hidden biological attributes. We use the vast prior knowledge regarding spatial zonation in hepatocytes to uncover the temporal trajectory in the data. At last, we present SiFT’s applicability to the case-control setting. In the context of COVID-19, SiFT exposes disease-related signals and single-cell dynamics by filtering a corresponding healthy trajectory obtained by mapping to reference healthy samples.

In contrast to various latent space representation methods, SiFT performs the correction at the level of individual genes. In turn, similarly to other data correction methods, it modifies the gene expression so that data is no longer “properly” log-transformed, a property that is expected in certain downstream analysis tasks. With that, this correction allows for obtaining biologically relevant insights on the gene level.

Here we exemplified the diverse range of applications of SiFT and showcased the potential that *filtering* has for understanding the different biological signals encoded in single-cell data. We further envision that with the ongoing increase in single-cell analysis tools along with the advance in multi-modal assays, SiFT will serve as a basic analysis tool revealing hidden, more complex structures in the data. We have made SiFT available as an open-source python package along with documentation and tutorials and ensured it can efficiently scale to the ever-increasing sizes of single-cell datasets.

## Methods

### The SiFT algorithm

The aim of SiFT is to expose hidden biological signals in an input count matrix. Given an expression matrix along with a mapping of the genes to a specific *signal*, SiFT computes the cell-cell similarity kernel based on this mapping, the projection of the data onto the *signal*, and the filtered expression matrix.

The input to SiFT includes a cell (*N*) by features (*G*) matrix 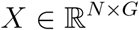 and a mapping of the cells, *T*. For best performance, we recommend that the matrix *X* will contain pre-processed normalized gene expression. In addition, SiFT can also be applied to any other representation of the cells, e.g. latent space or a subset of the genes.

The mapping, *T*, is assumed to capture a specific biological attribute or representation of the cells and can be of any type, e.g. stochastic or deterministic, continuous or discrete, uni- or multi-variate. The diversity of types of mappings that are supported includes, for example, donor age (deterministic, discrete, univariate), pseudotime reconstruction (deterministic, continuous, univariate), and a latent space representation of the data (PCA, scVI, etc.) (deterministic, discrete, multivariate). Alternatively, a set of process-specific marker genes (e.g. cell cycle marker genes), can be considered as a type of mapping and used to define distances between the cells. The type of mapping provided dictates the cell-cell similarity kernel that can be used.

The SiFT procedure comprises three main steps:

1. *Compute cell-cell similarity kernel (K*): In the first step, SiFT computes a cell-cell similarity kernel, *K* ∈ [0,1]^*N×N*^. The kernel is a row stochastic matrix, e.g. rows sum to one hence defining a proper distribution. The details of the kernel construction depend on the specific kernel choice (see below). In all cases the user can choose the subset of *source* and *target* cells, that is the cells for which the kernel is computed (*source*) and the cells the similarity is evaluated with respect to (*target*). The user can also provide a pre-computed kernel.
2. *Project the data (P*): obtain a projection of the data on the supervised *signal*, 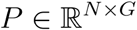,

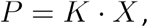

the stochasticity of *K* guarantees that *P* is of the same order of magnitude as *X*.
3. *Filter the data* (SiFTer, 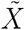): deduce the projection, *P* from the original count matrix *X* to obtain a *filtered* expression, 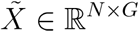,

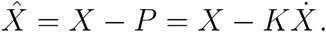

### The kernels

The kernel, *K* is supposed to capture the cell-cell similarity with respect to the *signal* we wish to SiFT. *K* is a stochastic matrix such that the *i*th row is a probability distribution denoting the similarity of cell *i* to all of the observed cells. Broadly speaking, the kernels can be divided into three sub-classes, *mapping, k-nearest-neighbor (knn*), and *distance* kernel, differing in the type of prior knowledge or assumptions they require. Beyond the implemented kernels the user can provide a pre-computed kernel.

All kernels can be refined by restricting the *source* and/or *target* space. The *source* space relates to the cells whose expression we are interested in filtering. The *target* space is the cells over which similarity is assessed. This is done by specifying the sub-group of cells (e.g. only healthy cells in a disease-control experiment) of interest.

#### Mapping kernel

The basis of the mapping kernels is a stochastic or deterministic association of the cells (*N*) to a low-dimensional domain. The mapping, *T* can be of any type, e.g. stochastic or deterministic, continuous or discrete, uni- or multi-variate. In the case of a continuous variable a binning value *M* is required, denoting the number of bins used to construct a binned representation. The mapping is represented by 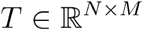. Given *T* we construct two sets of probability distributions:

1. *p_l_* (*μ* |*c_i_*): the probability of a label *μ* given a cell *i*, normalizing *T* across possible labels.
2. *p_c_* (*c_i_* |*μ*): the probability of observing a cell *i* given a label *μ*, normalizing *T* across cells.

The cell-cell similarity kernel, *K* ∈ [0, 1]^*N×N*^, is defined as the multiplication between the two probabilities, *p_l_* and *p_c_*, summing out the dimension of the embedded signal, *M*. Thus, each row in *K* induces a normalized distribution, *p_c_i__* (*c_j_*), defined for cell *i* with respect to all cells in the dataset

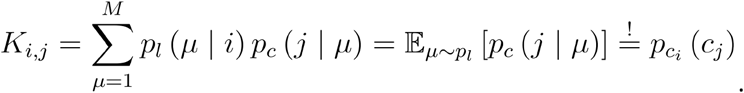

#### K-nearest-neighbor kernel

The mapping 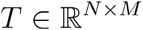 is used to construct a neighborhood graph over the cells by finding the *k* nearest neighbors for each cell. Following Scanpy’s^18^ defaults, the neighbor sets are merged and processed via the UMAP algorithm^20^. We normalize (across the rows) the resulting weighted adjacency matrix of the neighborhood graph of the cells (termed the connectivities) to obtain the final kernel object, *K*.

#### Distance kernel

A distance kernel requires that the provided mapping, 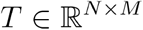, which defines a joint space representation of the cells, will be equipped with a distance metric, 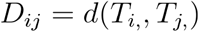 (e.g. the Euclidean distance). With this, we define the cell-cell similarity kernel as

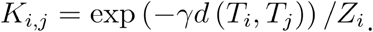

Here *γ* is a smoothing parameter that controls the effective radii of cells (distances) that have a non-negligible influence and *Z* is a normalization constant, defined for each row. If *d*(·,·) is chosen to be the Euclidean distance, the above denotes the radial basis function (RBF) kernel, a popular kernel function used in various kernelized learning algorithms which can be interpreted as a similarity measure ^38^. Similarly, if *d*(·,·) is chosen to be the Manhattan distance, then *K* is the Laplacian kernel.

### The stochastic interpretation of the SiFT algorithm and output

Looking into the mathematical derivation of the SiFT procedure we can expose natural stochastic properties which provide a better understanding of the SiFTed output. Recall that the input mapping, *T*, is used to define a row stochastic kernel, *K* ∈ [0, 1]^*N×N*^. That is 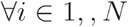 the *i*th row of *K* is a probability distribution for cell *i* with respect to remaining cells, we denote this distribution as *p_c_i__*(*c_j_*).

Now, an entry in the *projected data P_ik_* (for cell *i* and gene *k* in *P* = *K* · *X*, step 2 in the SiFT procedure), can be read as

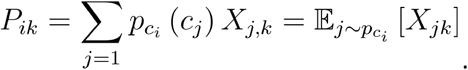

With this, for each cell, the projection 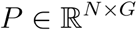 can be understood as the mean expression of genes according to the cell-cell similarity distribution. At the final step *P* is deduced from the expression, so we obtain

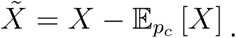

Thus, the filtered expression stands for the deviation of the cells’ expression with respect to the expected value in their *neighboring* cells. This implies that the resulting matrix can contain negative values, which translate to genes (or features) that are below the average expression. Generally speaking, we are only interested in the relations and distances between the cells in the new, filtered space, and not the absolute counts, hence the existence of negative values do not pose a problem. With that, Realizing that certain analysis methods expect as input a positive count matrix, we suggest *correcting* the filtered expression matrix by adding a pseudo count following the global minima of the data (so that the corrected filtered minima would be zero), ensuring positivity and preserving the topology of the data.

#### Runtime considerations

SiFT uses pyKeops, a python package allowing for Kernel Operations on the GPU without memory overflows^39^ as a backend for matrix computations. The implementation supports pyKeops pytorch and numpy backends and hence does not enforce pytorch dependency. As we show in Figure 2(c), this implies that SiFT can scale to large datasets without harming runtime performance. Furthermore, for datasets too large to be handled by a single GPU operation, instead of falling back to CPU computation we implement an automatic batching procedure, which retains speed performance.

### Datasets

#### Drosophila wing disc myoblast cells

We obtained the dataset of a temporal cell atlas of the Drosophila wing disc from two developmental time points collected by^17^ and using the processed data published by^9^, available at myoblasts.h5ad. The data contains two biological replicates were obtained at each time point that, after filtering for low-quality cells, generated data from 6922 and 7091 cells in the 96 hr samples and 7453 and 5550 cells in the 120 hr samples.

In order to quantitatively assess our performance in removing unwanted variation with respect to these attributes, we turned to classify the cells into sex and cell cycle phase categories. The processed dataset was lacking sex and cell cycle labels however there are known marker genes for both. For sex labels, we follow the procedure suggested in the original study in which the data was presented^17^. The classification relies on the expression levels of the dosage compensation complex genes *lncRNA:roX1* and *lncRNA:roX2*. For both genes, we computed the density over the log-normalized counts and identified the first local minima as a threshold (Figure 2, Supplementary Figure 1). Cells that were above the threshold for either *lncRNA:roX1* or *lncRNA:roX2* were classified as male; otherwise, they were classified as female.

Next, to obtain cell cycle phase categories we applied Scanpy’s “scanpy.tl.score_genes_cell_cycle” based on the expression of known Drosophila cell-cycle genes from Tinyatlas at Github^40,41^ (Figure 2).

##### Methods application

To apply SiFT we defined a *knn* kernel using the set of cell cycle and sex marker genes (see Supplementary Table 1). For the regress_out setting, Scanpy’s “scanpy.pp.regress_out” function was used by setting all marker genes as regression keys. For scVI we followed the reproducibility notebook, [scvi_covariates.ipynb](https://yoseflab.github.io/scvi-tools-reproducibility/scvi_covariates/) using only the set of cell cycle and sex marker genes as continuous covariates.

##### scVI corrected expression

scVI provides the user with the option to impute normalized corrected counts through “get_normalized_expression()”. We use this function followed by “sc.pp.log1p()” to obtain the corrected counts used for HVG evaluation. It is important to note that scVI is not designed for this task precisely, hence, we evaluated its performance on the latent embedding when possible^9^.

##### Quantitative evaluation

To evaluate the removal of cell cycle and sex *signals* we use the graph iLISI score as defined in^42^, using the corrected expression for SiFT and regress_out and over the latent embedding for scVI. We evaluated the graph iLISI score independently with respect to the “cell cycle phase label” and the combined ”cell cycle phase and sex label”.

To assess the perseverance of the biological signal of interest we use the set of genes of interest suggested by^9^ which is based on marker genes presented in^17,19^ (see Supplementary Table 1). Here we look at the intersection of this set with the set of highly variable genes (HVGs). HVGs identifies using Scanpy’s “scanpy.pp.highly_variable_genes” with n_top_genes=500 and flavor=”cell_ranger”. Since this can not be done over the latent space we resort to using the imputed gene expression provided by scVI.

##### Batch integration

For additional evaluation regarding full integration performance following SiFT and regress_out, we applied batch integration methods. We used bbknn^7^ and Harmony^8^. Both were run using Scanpy’s functions with default parameters passing the batch key for integration. For comparison, we use scVI with the batch label as a batch key along with the continuous covariates (scVI (covariates & batch)). Here, when computing the graph iLISI score, the latent embedding was used for all methods as neither bbknn nor Harmony provides a correction to the gene expression.

#### Heart Cell Atlas dataset

the Heart Cell Atlas dataset was downloaded from https://www.heartcellatlas.org and contains a total of 486,134 cells.

##### Benchmarks data

For the runtime analysis we followed the procedure suggested by Gayoso et al.^9^. Using the Heart Cell Atlas dataset which contains 486,134 cells, we created 8 datasets of increasing size by subsampling 5,000, 10,000, 20,000, 40,000, 80,000, 160,000, 320,000, and 486,134 cells. For each dataset the top 4,000 genes were selected via “scanpy.pp.highly_variable_genes”, with parameter flavor=“seurat_v3”. Next, we generated 8 random covariates by sampling from a standard normal distribution and used them along with the *percent_mito* and *percent_ribo* fields as continuous covariates, defining a total of 10 continuous covariates.

##### Methods runtime analysis

Performed on NVIDIA RTX A5000 GPU. For SiFT runtime, we report the runtime of initialization of the *rbf* kernel and running the filtering procedure. For scVI runtime, we report the runtime of the train function with the parameters used in^9^:

early_stopping=True, early_stopping_patience=45, max_epochs=10000, batch_size=1024, limit_train_batches=20, train_size=0.9 if n_cells < 200000 and train_size=1-(20000/n_cells) otherwise

For the regress_out baseline, we tracked the runtime of the regress_out function for the above continuous covariates.

#### Virtual tumor data

The simulated dataset was downloaded from Cyclum’s repository (https://github.com/KChen-lab/Cyclum/tree/master/old-version/data/mESC). Details regarding the simulations of the virtual tumor data can be found in the original publication^10^. This data contains a total of 279 cells, 168 in belong to the tumor “intact” and 111 to the “perturbed”. Cell’s are given with ground truth labels regarding the cell cycle phase.

##### The kernels

We consider different mappings representing the cell cycle signal. In total six different SiFT kernels are used for comparison:

- *ground truth labels*: A *mapping kernel* where the discrete cell labels are given by the ground truth cell cycle phase labels.
- *cell cycle genes*: A *knn kernel* where cell neighbors are computed based on the expression of the set of cell cycle genes reported in^43^.
- *Cyclum pseudotime*: we use Cyclum’s pseudotime mapping of the cells in order to define three kernels:
  - *Cyclum, binned (n=12*): A *mapping kernel* where the discrete cell labels are obtained by binning the pseudotime to 12 bins.
  - *Cyclum, binned (n=3*): Same as above with three bins.
  - *Cyclum, distance*: A *distance kernel* where distances are computed over the mapping of the cells to the unit circle using the pseudotime, (*x, y*) = (cos(*pseudotime*), sin(*pseudotime*)).
- *Seurat*: A *mapping kernel* where the discrete cell labels are given by Seurat’s cell cycle phase prediction.

##### Cell cycle removal methods

- Cyclum: we followed the steps in the provided example by the authors, example_mESC_simulated.ipynb. As the example is provided using an older version of Cyclum, we modified the parameters in the new implementation to correspond to the reported setup run in^10^.
- Seurat: we followed the steps suggested in the vignette cell_cycle_vignette.html.
- ccRemover: we used the ccRemover method as in the tutorial ccRemover_tutorial.html.

##### Evaluation metrics

Evaluation metrics and their definitions were taken from^23^. We used the complementary python package scib (https://scib.readthedocs.io) which provides an implementation of all metrics. For the “cell cycle removal” score we reported the mean of *ASW_label/batch, PCR_batch, cell_cycle_conservation*, and *iLISI_graph*. For the “bio conservation” score we reported the mean of *NMI_cluster/label, ARI_cluster/label, ASW_label, isolated_label_F1, isolated_label_silhouette*, and *cLISI_graph*. All methods were run with default parameters.

#### Mammalian liver dataset

ScRNA-seq data was downloaded from GEO, accession code GSE145197. For the data pre-processing procedure we followed the pipeline described in the original publication^24^ and given in the GitHub repository https://github.com/naef-lab/Circadian-zonation. After pre-processing the data contained 11,491 cells from 3 biological repetitions for 4 different time points (time point 0: 3563 cells, time point 6: 2791 cells, time point 12: 2919 cells, time point 18: 2218 cells).

##### novoSpaRc for spatial mapping

We used novoSpaRc^2,3^ to recover the spatial signal in the data, and obtain a probabilistic mapping of cells to the eight liver zonation layers. We performed mapping using 15 spatial marker genes (reported in^24^) and ran the novoSpaRc algorithm with *α* = 0.5 and *ϵ* = 1*e* – 1. We used the probabilistic mapping to apply SiFT (Supplementary Figure 5).

#### COVID-19

The data object, as an h5ad file, was downloaded from https://covid19cellatlas.org/, haniffa21.processed.h5ad. Cell type labels and metadata were based on the fields reported in the given file. We considered COVID-19 samples (*n_disease_* = 527, 286) and healthy controls (*n_healthy_* = 97, 039).

##### Distance kernel construction

To filter the healthy trajectory we define a *knn kernel n_disease_* × *n_healthy_* that captures the similarity of the disease cells to the reference healthy cells. To evaluate the similarity (distances) between the disease and healthy cells we consider distances in the harmonized PCA space (reported by^27^).

##### cluster_purity score

We define a cluster purity score as a measure to distinguish between indicative and non-indicative clusters with respect to a given label *y* (here we consider the cell type). Formally, for a given cluster of size *n*, we find the most frequent label, *y*, with *m* cells. Now, we say that a cluster is indicative if

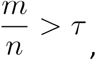

for a given threshold value *τ*. This measure quantifies the homogeneity of the cluster.

## Data availability

The datasets analyzed in the current study are available at

- Drosophila wing development: https://figshare.com/articles/dataset/scvi-tools-reproducibility_processed_data/14374574/1?file=27458846
- Heart Cell Atlas: https://cellgeni.cog.sanger.ac.uk/heartcellatlas/data/global_raw.h5ad
- Virtual tumor: https://github.com/KChen-lab/Cyclum/tree/master/old-version/data/mESC
- Mammalian liver: https://www.ncbi.nlm.nih.gov/geo/query/acc.cgi?acc=GSE145197
- COVID-19: https://covid19.cog.sanger.ac.uk/submissions/release1/haniffa21.processed.h5ad

## Code availability

Software is available at https://github.com/nitzanlab/sift-sc and documentation at https://sift-sc.readthedocs.io. The code to reproduce the results is available at https://github.com/nitzanlab/sift-sc-reproducibility.

## Acknowledgments

We thank Michal Klein for his thoughtful review and assistance in the software development, and Klaas Mulder and Sabine Tanis, for ongoing discussions regarding the presented framework. We would further like to thank Kristen Koenig and Inbal Avraham-Davidi for stimulating biological discussions and Aleksandrina Goeva for helpful insight. We acknowledge Noa Moriel, Jonathan Karin, and all members of the Nitzan lab for general feedback.

