## Supplementary information for "Uncovering hidden biological processes by probabilistic filtering of single-cell data"

Zoe Piran[1] and Mor Nitzan[1,2,3*]

[1]School of Computer Science and Engineering, The Hebrew University, Jerusalem, Israel.
[2]Racah Institute of Physics, The Hebrew University, Jerusalem, Israel.
[3]Faculty of Medicine, The Hebrew University, Jerusalem, Israel.

Supplementary Figures

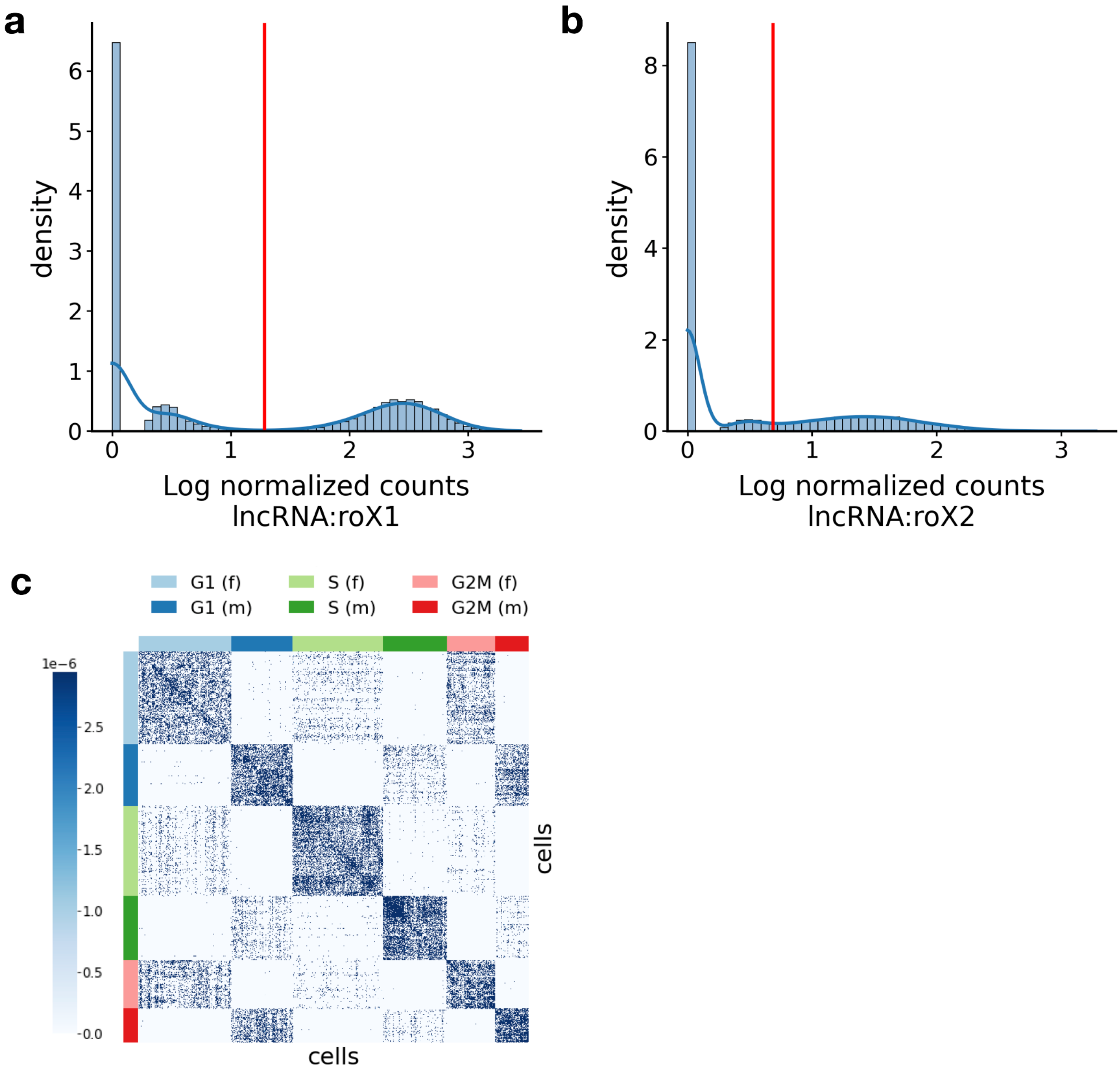


**Supplementary Figure 1:** Pre-processing for the removal of unwanted sex and cell cycle effects from single-cell transcriptomics of the Drosophila wing disc development. **(a)-(b)** Probability histogram plots of the log normalized expression counts for lncRNA:roX1 (a) and lncRNA:roX2 (b) within all cells. Density curves for the data are shown in blue. Red lines are drawn on the first local minima within the density of the data and serve as a cutoff for classifying cells as having a high or low expression of either gene. Cells with high expression of either lncRNA:roX1 or lncRNA:roX2 were classified as male-originating; otherwise, cells were designated as female-originating **(c)** The SiFT kernel, a *knn kernel* based on the graph connectivity matrix based on the set of sex and cell cycle genes. Cells are ordered according to the combined label of sex and cell cycle phase.

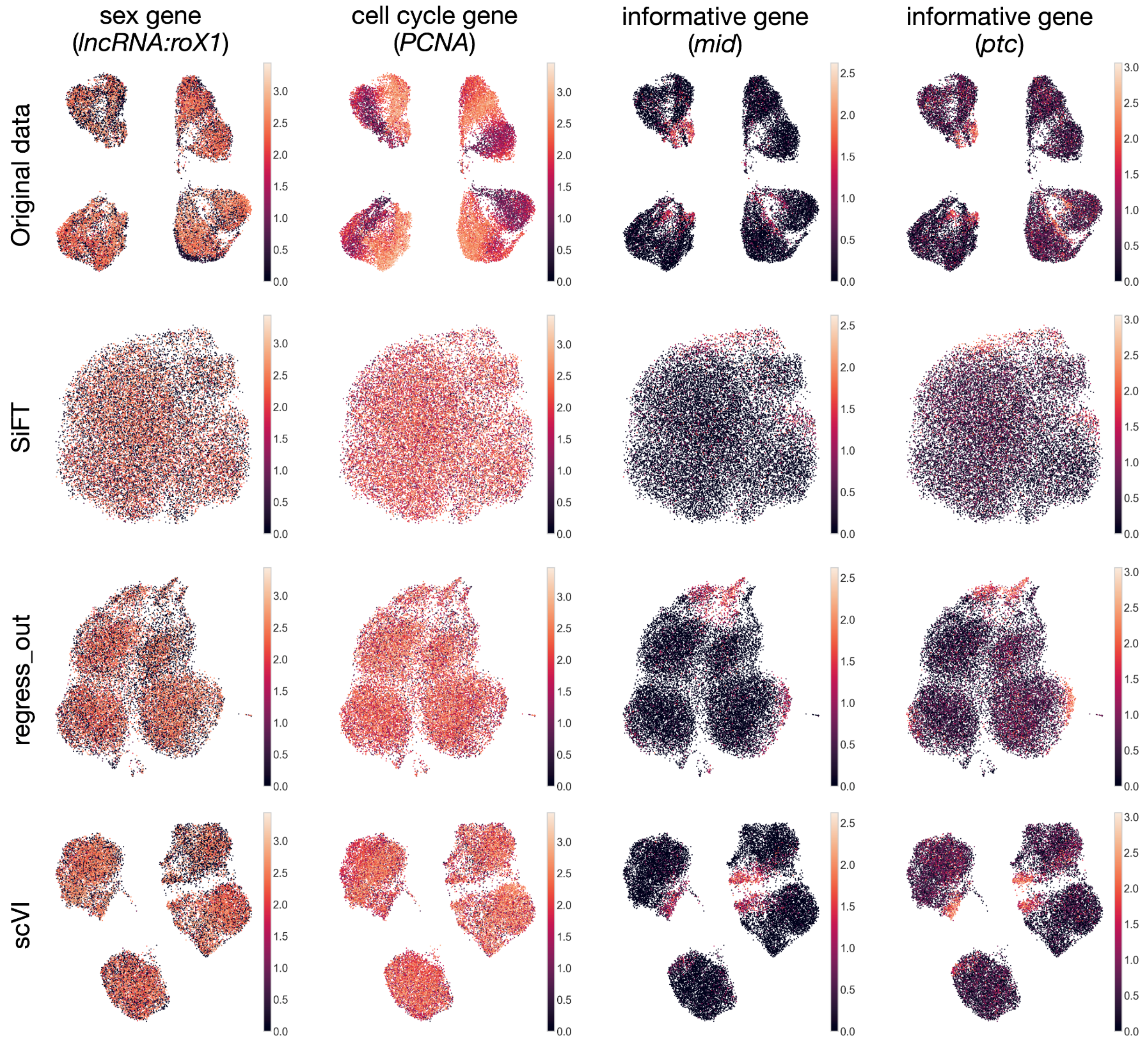


**Supplementary Figure 2**: removal of unwanted sex and cell cycle effects from single-cell transcriptomics of the Drosophila wing disc development. UMAP embeddings following different data correction procedures (rows) and colored by genes representing different data features (columns). Rows (top to bottom) show uncorrected data (Original data), SiFT filtered (SiFT), Scanpy's "scanpy.pp.regress_out()" (regress_out), and scVI latent space with continuous covariates correction (scVI). Columns (left to right) *lncRNA:roX1* (sex gene), *PCNA* (cell cycle gene), *mid* and *ptc* (informative genes reported by[1]).

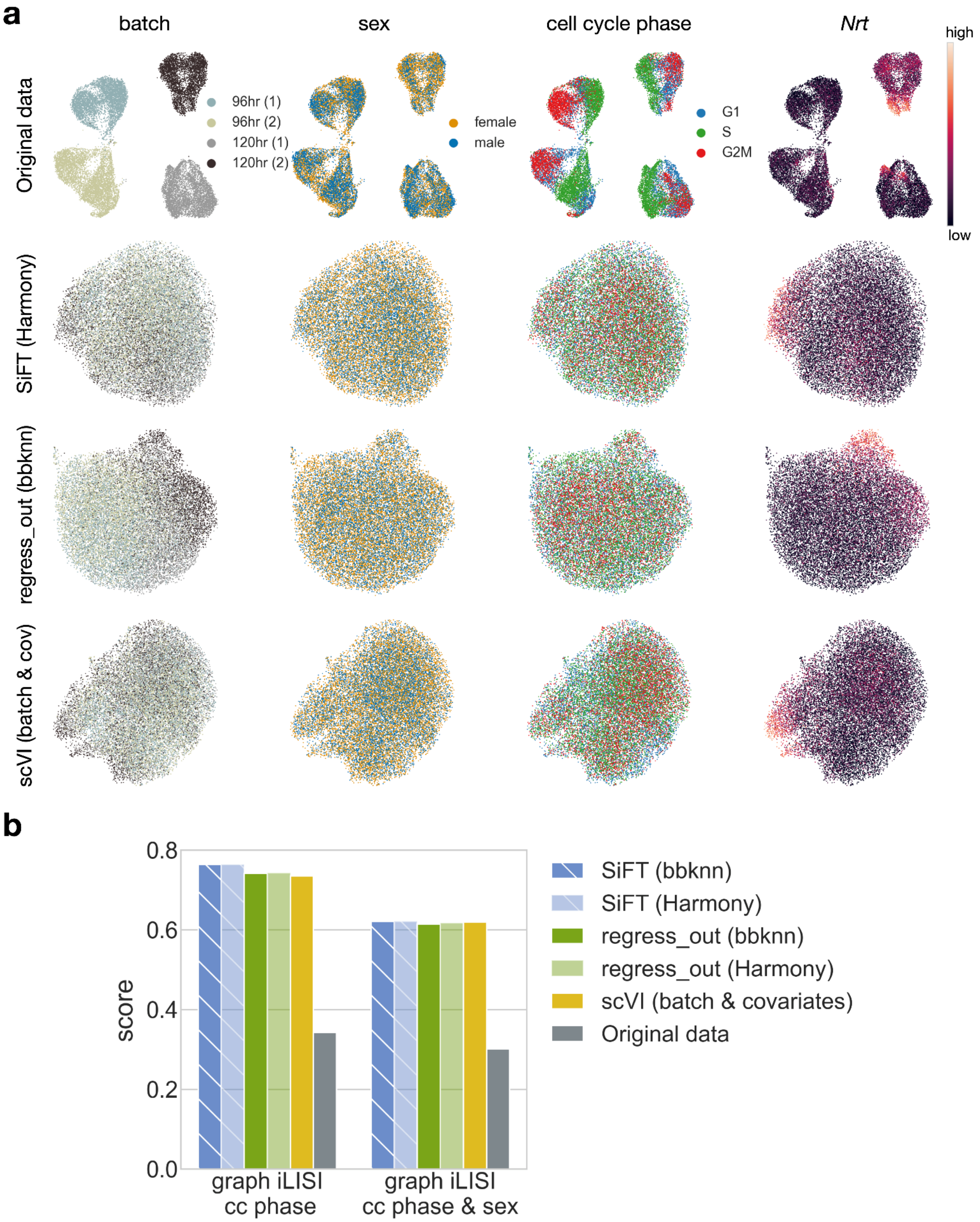


**Supplementary Figure 3**: Batch integration along with the removal of variation from single-cell transcriptomics of the Drosophila wing disc development[1]). **(a)** UMAP embeddings following different data correction procedures (rows) and colored by different covariates of unwanted sources of variation (columns). Rows (top to bottom) show uncorrected data (Original data), SiFT filtered followed by Harmony integration (SiFT (harmony)), regress_out followed by bbknn (regress_out (bbknn)), and scVI latent space with batch and continuous covariates (scVI (batch & cov.)). Columns (left to right) show the batch label, sex label, cell cycle phase, and cell cycle and sex (see Methods). **(b)** The graph iLISI score each data correction procedure obtained for the different covariate labels.

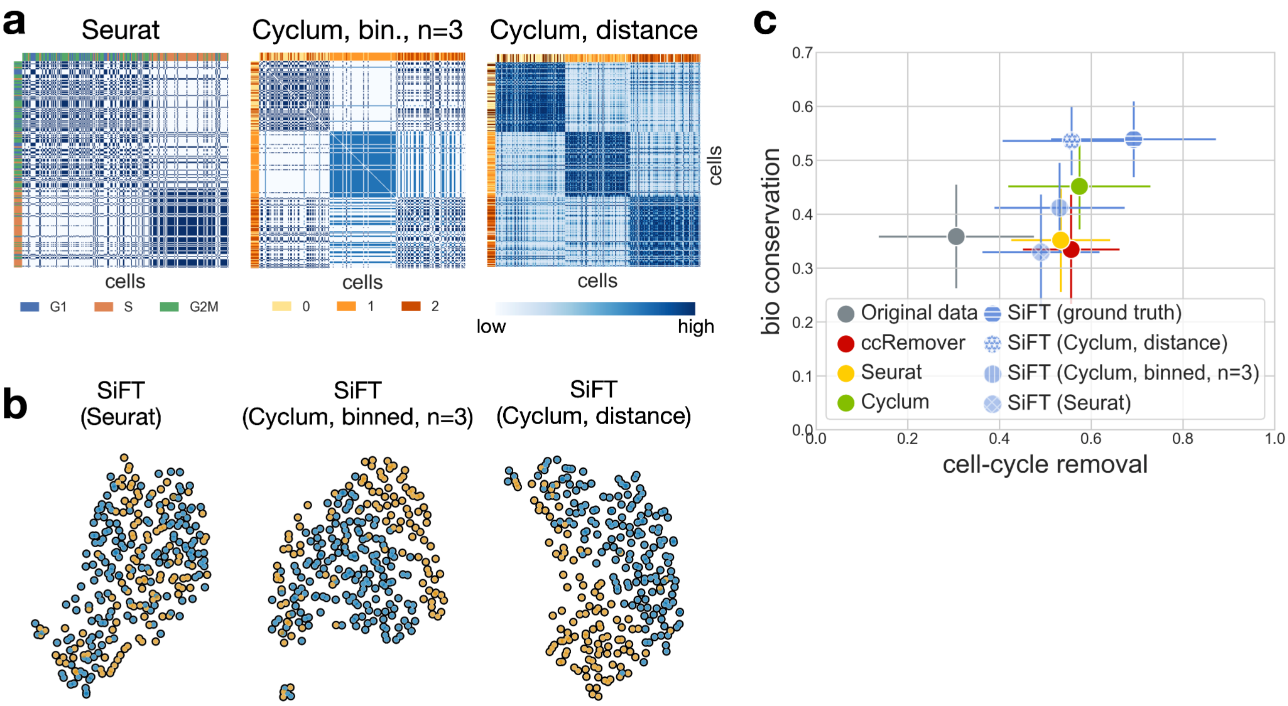


**Supplementary Figure 4:** Filtering the cell-cycle effects from the virtual tumor data consisting of two subclones. **(a)** Different cell-cell similarity kernels are defined by SiFT. Cells are ordered according to the ground truth cell cycle stage. (left) a *mapping kernel* based on Seurat cell cycle stage classification. The row(col) colors depict the Seurat inferred cell cycle stage. (center) a *mapping kernel* based on binning of the Cyclum pseudotime ($$n$$, number of bins, $$n=3$$), the row(col) colors depicts the cells' bin. (right) a *distance kernel* distances defined over the Cyclum pseudotime prediction. **(b)** UMAP of the filtered data colored by the sub-clone identity. (left) Seurat cell cycle stage prediction (center) Cyclum binning ($$n=3$$) (right) distance in Cyclum pseudotime **(c)** Scatter plot of the mean overall *bio conservation score* against mean overall *cell cycle removal scores* using the metrics defined in[2] (see Methods). The error bars indicate the standard mean error.

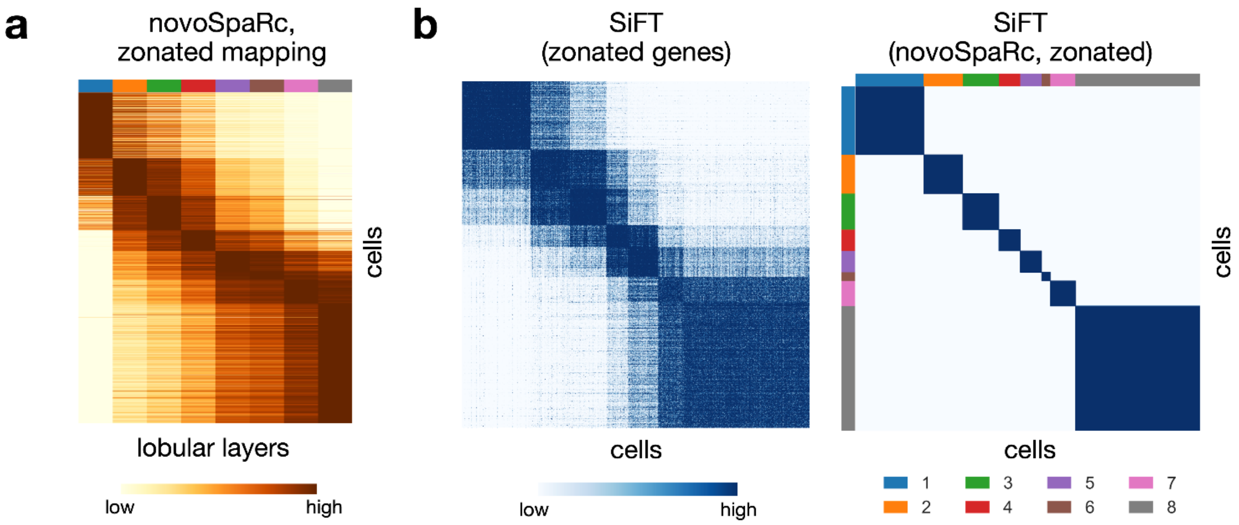


**Supplementary Figure 5:** Enhancing circadian clock signal in the mammalian liver. **(a)** The mapping of the mammalian liver to eight lobular layers as obtained by novoSpaRc[3,4]. **(b)** The SiFT kernels used for filtering.

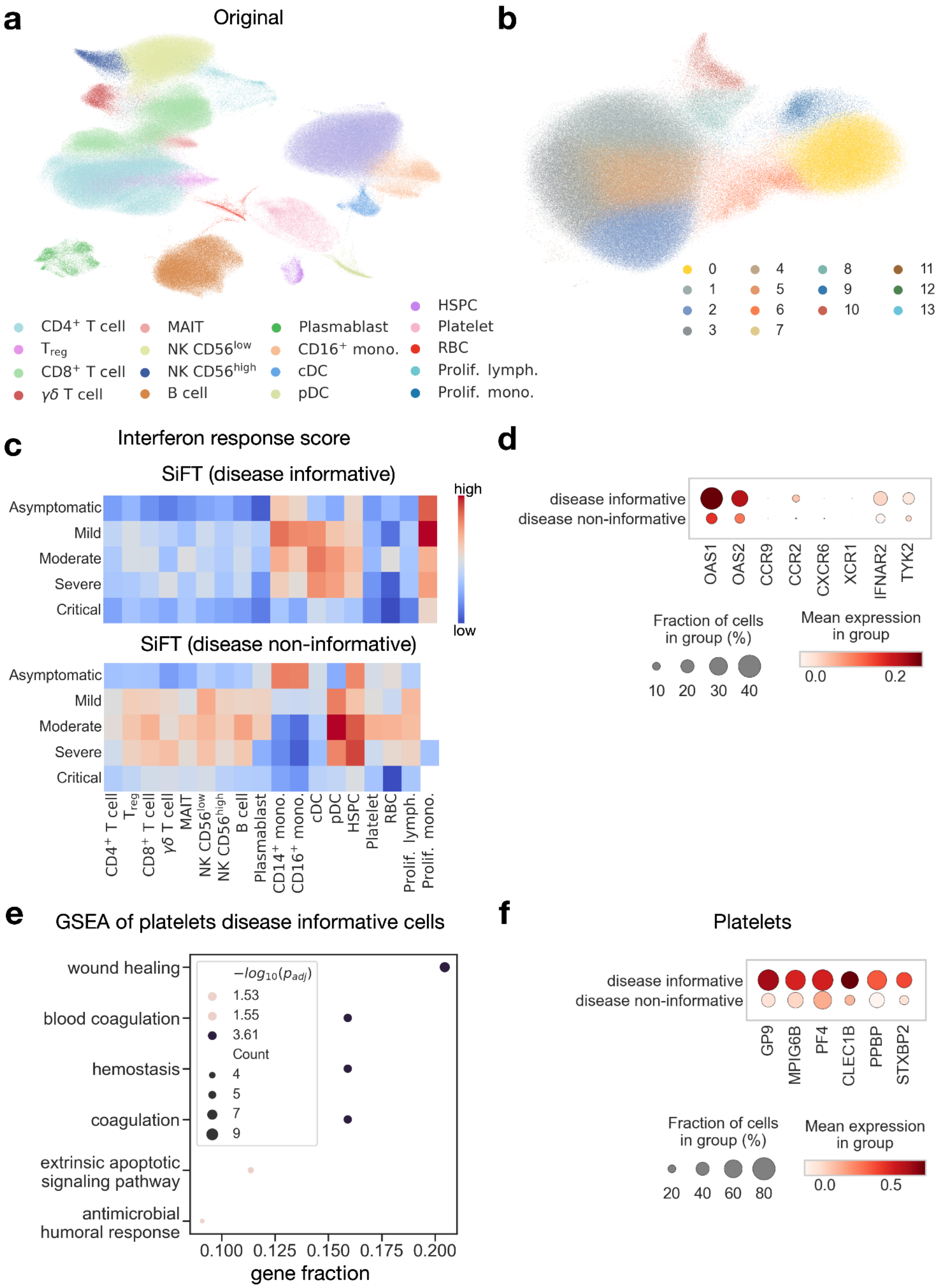


**Supplementary Figure 6:** Revealing the disease signal in COVID-19 dataset. **(a)** UMAP visualizations of cells in the original data colored by reported cell types. **(b)** UMAP visualizations of cells in the data after applying SiFT, colored by Leiden clustering of the SiFTed data. **(c)** Enrichment of interferon response of each cell state separated by disease severity. Shown for SiFTed disease informative cells (top) and SiFTed disease non-informative cells (bottom). IFN response was calculated using a published gene list (GO:0034340). **(d)** Dot plot of gene expression where the color is scaled by mean expression and the dot size is proportional to the percent of the population expressing the gene considering genes associated with COVID-19 identified in recent GWAS studies[5,6] **(e)** GSEA of differentially expressed genes in the $Platelets$ disease informative cells (compared to non-informative, using top 50 genes). Size of circles indicate the number of genes. Color indicates the magnitude of $-\log_{10}(p_{adj})$. **(f)** Dot plot of gene expression of platelets activation markers in the $Platelets$ disease informative and non-informative cells. Markers taken from[7].

### Supplementary Tables

| Sex Genes | Cell cycle genes | Genes of interest |
|---|---|---|
| lncRNA:roX1 | PCNA | Argk |
| lncRNA:roX2 | dnk | Nrt |
| Sxl | RnrS | Ten-a |
| msl-2 | RnrL | Ten-m |
| | Claspin | wb |
| | Mcm5 | Act57B |
| | Caf1-180 | drl |
| | RPA2 | mid |
| | HipHop | nemy |
| | stg | lms |
| | Mcm6 | CG11835 |
| | dup | Gyg |
| | WRNexo | ara |
| | Mcm7 | tok |
| | dpa | kirre |
| | CG10336 | NK7.1 |
| | Mcm3 | fj |
| | Mcm2 | beat-IIIc |
| | RpA-70 | CG33993 |
| | Chrac-14 | dpr16 |
| | CG13690 | CG15529 |
| | RPA3 | CG9593 |
| | asf1 | beat-IIb |
| | CDC45L | robo2 |

| | | |
|---|---|---|
| | DNApol-alpha73 | Ama |
| | CycE | fz2 |
| | DNApol-alpha50 | elB |
| | Kmn1 | noc |
| | Lam | nkd |
| | Nph | fng |
| | msd5 | vg |
| | msd1 | |
| | ctp | |
| | Set | |
| | scra | |
| | Chrac-16 | |
| | ncd | |
| | Ote | |
| | pzg | |
| | HDAC1 | |
| | nesd | |
| | tum | |
| | CG8173 | |
| | aurB | |
| | feo | |
| | pav | |
| | CG6767 | |
| | sip2 | |
| | Det | |
| | Cks30A | |
| | CycB | |
| | B52 | |

**Table 1**. Gene sets for multiple covariates Drosophila analysis (based on[8]).

| Spatial Genes | Temporal genes |
|---|---|
| Glul | Bmal1 (Arntl) |
| Ass1 | Clock |
| Asl | Npas2 |
| Cyp2f2 | Nr1d1 |
| Cyp1a2 | Nr1d2 |
| Pck1 | Per1 |
| Cyp2e1 | Per2 |
| Cdh2 | Cry1 |
| Cdh1 | Cry2 |
| Cyp7a1 | Dbp |
| Acly | Tef |
| Alb | Hlf |
| Oat | Elov3 |
| Aldob | Rora |
| Cps1 | Rorc |

**Table 2**. Spatial and temporal gene sets used in the analysis of the liver dataset (based on[9]).
